# Interstitial pneumonia via the oropharyngeal route of infection with *Encephalitozoon cuniculi* Oropharyngeal infection with *Encephalitozoon cuniculi*

**DOI:** 10.1101/2024.04.04.588046

**Authors:** Mariana Souza Santos, Rodrigo A. da Silva, Lucas Nunes de Souza, Nicolli R. Victorino, Rayane C.B. Pereira, José Guilherme Xavier, Ronalda A. da Silva, Elizabeth C. Pérez, Anuska M. Alvares-Saraiva, Maria Anete Lallo

## Abstract

Microsporidia causes opportunistic infections in immunosuppressed individuals. Mammals shed these spores of fungi in feces, urine, or respiratory secretions, which could contaminate water and food, thereby reaching the human body and causing infection. The oral route is the most common route of infection, although experiments have demonstrated that intraperitoneal and intravenous routes may also spread infection. Respiratory tract infection, although considered to be possible, has not been reported to date. The present study, therefore, aimed to demonstrate infection with the microsporidia of *Encephalitozoon cuniculi* via the oropharyngeal route as a model for opportunistic pneumonia. The objectives were the study of opportunistic pneumonia in general while also confirming transmission via the respiratory route to expand the understanding of the epidemiology of zoonoses. C57BL mice, both male and female, up to 12 weeks of age, and free of specific pathogens (SPF) were inoculated (Infected group) or not infected (Non-Infected group) with 1 × 10^7^ spores of *E. cuniculi* via the oropharyngeal route. The animals immunosuppressed with cyclophosphamide (Cy) were infected (Cy-Infected group) or not infected (Cy-Uninfected group), and then assessed for the influence of immunosuppression on infection. In both groups, animals inoculated with 0.9% saline solution via the oropharyngeal route served as controls (Sham and Cy-Sham groups, respectively). After 14 days of infection, the lungs of all animals were retrieved for histopathological analysis, phenotyping of lung inflammatory cells using flow cytometry, measurement of fungal load using qPCR, and measurement of the serum levels of Th1, Th2, and Th17 cytokines. The results revealed that the infected animals developed interstitial pneumonia characterized by perialveolar inflammatory infiltrative lesions with a predominance of lymphocytes and plasma cells. However, the fungal load and the extent of the inflammatory infiltrate were relatively lower in the Cy-Infected group, with a predominance of the CD8^+^ T lymphocyte population in the lungs compared to Infected group. In the Infected group not treated with Cy, an increase in the population of alveolar macrophages (F4/80^+^CD11b^-^SiglecF^+^) was noted, along with higher fungal load and inflammatory infiltrate in the lungs, indicating a further pronounced *Encephalitozoon* pneumonia compared to the immunosuppressed animals. The infected groups presented Th1 cytokine profiles, with the Cy-Infected group exhibiting increased Th17 levels. Collectively, these results demonstrated that oropharyngeal infection promoted pneumonia caused by *E. cuniculi* in mice treated or not with cyclophosphamide, with greater severity occurring in the non-immunosuppressed mice, thereby establishing this model as a suitable one for interstitial pneumonia via aspiration.

**Author Summary:** Opportunistic fungi infection in immunosuppressed individuals may lead to pneumonia that is lethal. The airways are the most common routes for these infections. The opportunistic fungal species *Encephalitozoon cuniculi* is associated with pneumonia, although this species has not been reported to cause airway infection to date. The present study aimed to demonstrate that the oropharyngeal route, which is a frequent route of infection in patients with reflux, could lead to severe pneumonia in the mice without immunosuppression associated with a low CD8^+^ T lymphocyte response.

## Introdution

Microsporidia are obligatory intracellular parasitic fungi belonging to phylum Microsporidia, which form spores of sizes ranging from 1 to 12 µm, each spore having a thick wall and a spiral polar tubule, and are responsible for cellular infection, a unique characteristic of this phylum [1]. These primitive eukaryotic pathogens, devoid of mitochondria, are responsible for opportunistic infections, i.e., the pathogens establish within the host by exploiting the compromised immune system of the host [2]. Microsporidiosis in mammals is caused by 17 species, among which over 1,700 species have been identified to be associated with this group of fungi, particularly with *Enterocytozoon bieneusi*, *Encephalitozoon intestinalis*, *Encephalitozoon cuniculi*, and *Encephalitozoon hellem* [2, 3]. Intestinal and systemic infections are the most commonly reported ones, with the pathogenic aspects of these infections varying according to the species involved and the quality of immune response elicited in the host [4].

Opportunistic respiratory infections are an important factor for determining patient prognosis and also crucial contributors to increased length of stay at the hospital and drastically increased risk of death in immunocompromised patients [5]. Microorganisms may infect the lower respiratory tract in the following four ways: 1) aspiration of the contents from the oropharynx; 2) inhalation of contaminated aerosols; 3) spread of infection through contiguous regions; 4) hematogenous dissemination through extrapulmonary infectious regions (for example, in gastrointestinal tract infection). Among these four ways, two deserve particular attention: aspiration and hematogenous diffusion. A study reported that aspiration of microorganisms originating from the upper airways during sleep occurred in 45% of healthy patients and 70% of the patients with impaired perception, such as those with alcohol dependency, drug consumption, epilepsy, and usage of sedation [6]. The lung is the primary site of such infections, regardless of the infecting species, although immunosuppression does render the host susceptible to clinical pneumonia [5].

Microsporidian spores might be acquired by ingestion, inhalation, direct contact with the conjunctiva, contact with animals, or individual-to-individual transmission, ultimately causing infection [7]. In infections with microsporidia have been conducted experimentally in mice, and other rodents via oral, hematogenous, or intraperitoneal routes. In *E. cuniculi*, these routes of infection reportedly caused a disseminated disease with respiratory compromise, particularly in immunocompromised individuals [3, 8]. A study reporting the signs of symptomatic respiratory infection by *E. cuniculi* in kidney transplant recipients concluded that lifelong immunosuppression increases the risk of infection with respiratory microsporidiosis in these patients [9]. Moreover, while the cases of *E. cuniculi* pneumonia in immunocompromised patients, particularly the transplant recipients, are reported, infections with the respiratory route have not been elucidated to date. In this context, the present study aimed to evaluate infection with *E. cuniculi* via the oropharyngeal route in mice immunosuppressed with cyclophosphamide or those non-immunocompromised, as a model of infection caused by aspiration of oropharyngeal contents.

## Methods

### Ethics Statement

All experimental procedures were in accordance with the Brazilian law on the use of experimental animals, safety, and use of pathogenic agents. The study was approved by the Ethics Committee for Animals, Universidade Paulista (approval number: 1528190220, 04/02/2020).

### Animals

Specific-pathogen free (SPF) C57BL/6 male and female mice (n = 30), each 10 to 12-week-old, were procured from the “Centro de Desenvolvimento de Modelos Experimentais para Biologia e Medicina” at the Federal University of São Paulo, Brazil. The animals were housed in microisolators with controlled temperature (22 to 24 °C) and humidity (45% to 55%) under SPF conditions and a 12 h light and 12 h dark photocycle, throughout the experimental period, at the Animal Facility of Paulista University. The animals were provided with irradiation-sterilized pelleted food and autoclave-sterilized water ad libitum.

### E. cuniculi spores

*E. cuniculi* (genotype I) spores were purchased from Waterborne Inc. (New Orleans, LA, USA). The spores were grown in a rabbit kidney cell line (RK-13, ATCC CCL-37) using Roswell Park Memorial Institute (RPMI) medium supplemented with 10% fetal calf serum (FCS) and gentamicin at 37 °C under 5% CO_2_ atmosphere. The spores were then harvested from the tissue culture supernatants, washed thrice in phosphate-buffered saline (PBS), and finally counted using a hemocytometer.

### Cyclophosphamide treatment

Mice in the immunosuppressed group were treated with 200 mg/kg Cy via intraperitoneal injection once a week (Genuxal®, Asta Medica Oncologia, São Paulo, Brazil). This treatment was begun 14 days prior to pathogen inoculation followed by a weekly administration throughout the experimental period [10].

### Oropharyngeal inoculation in mice

The mice were anesthetized with sevoflurane via dispersion within a housing chamber until the animals lost movement and allowed the administration of the pathogen. Afterward, each animal was suspended, and its tongue was pulled laterally for administering the inoculum into the oropharynx using a micropipette. As an autonomous reflex action, the mouse inhaled the inoculated solution [11]. After inhalation, the tongue of the mouse was released, and the mouse was placed in a cage and monitored until it woke up from the effect of anesthesia. Later, the treatment mice were infected, through oropharyngeal aspiration, with 1 × 10^7^ spores of *E. cuniculi* dispersed in 50 µL of sterilized phosphate-buffered saline (PBS) solution. The Sham (control) group of mice, either treated or not treated with Cy, received 50 µL of sterilized 0.9% NaCl solution via the same route.

### Experimental design

The mice were subjected to two different protocols based on whether they had to be treated with Cy. Protocol 1 was applied to 3 groups of mice: Control group (n = 5) – untreated and uninfected mice; Infected group (n = 5) – mice infected with *E. cuniculi* spores via the oropharyngeal route; Sham group (n = 5) – mice that received 50 µL of sterilized the 0.9% NaCl solution. Protocol 2 was applied to the following groups: Cy-non-Infected (n = 5) – mice immunosuppressed with Cy and not inoculated with the pathogen; Cy-Infected (n = 5) – mice immunosuppressed with Cy and inoculated with the spores of *E. cuniculi*; Cy-Sham (n = 5) – mice immunosuppressed with Cy and inoculated with 50 µL the sterilized 0.9% NaCl solution. Protocol 2 was begun 14 days prior to infection (-–14 days), with the administration of a weekly dose of Cy as the immunosuppressive agent prior to the experimental infection. In the above 2 protocols, the course of infection was 14 days, after which the animals were euthanized and the tissue samples were collected for infection evaluation. The evaluations included the: histopathological analysis of the lungs and liver, phenotyping of the inflammatory cells in the lung, blood and bronchoalveolar lavage (BAL) determination, measurement of the levels of cytokines in the blood, and determination of fungal burden in the lung (Fig. 1).

**Figure 1.**
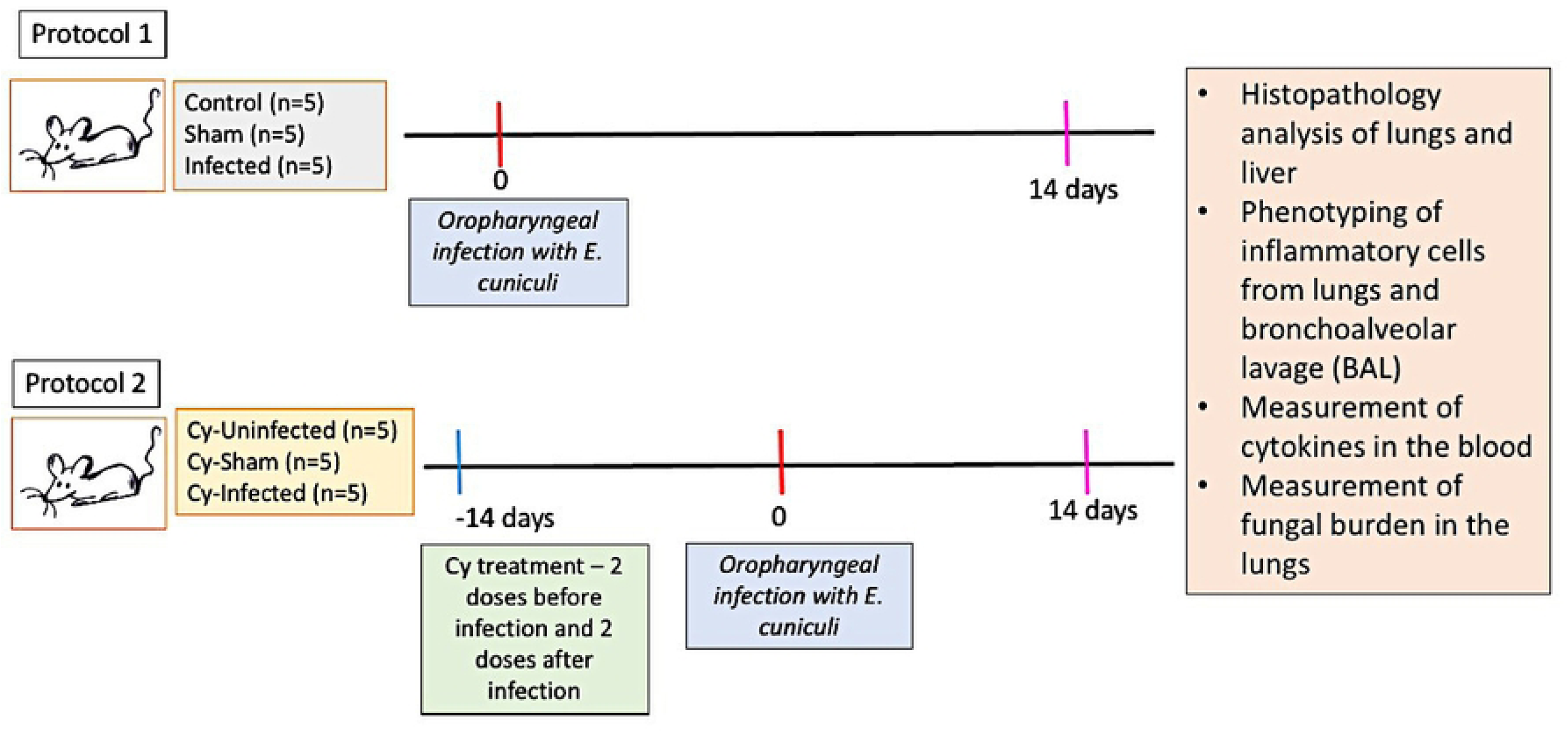
Experimental design. C57BL/6 mice were infected [or not infected, as required] with the spores of *E. cuniculi* followed by treatment [or no treatment, as required] with cyclophosphamide (Cy). After 14 days of infection, serum samples were collected to evaluate the levels of Th1, Th2, and Th17, histopathological analysis of the lungs and liver, phenotyping of the inflammatory cells in the lung, blood and bronchoalveolar lavage (BAL), measurement of the levels of cytokines in the blood, and determination of fungal burden in the lung.

### Necropsy and sample collection

After applying deep anesthesia using ketamine (100 mg/kg), xylazine (2.5 mg/kg), and acepromazine (2.5 mg/kg) intraperitoneally, blood samples were collected, via cardiac puncture, in a tube containing EDTA. The tubes with blood samples were then centrifuged to obtain the plasma samples, which were stored at –80 °C until use. Afterward, the thoracic cavity of each mouse was exposed, and bronchoalveolar lavage (BAL) was performed to isolate the trachea and fix it for the infusion of 1 mL of HBSS along with 2% FBS through aspiration. This procedure was conducted twice consecutively, and the collected material was centrifuged at 2000 rpm for 5 min. Finally, the sediment containing the cells was distributed in the wells of a V-bottom 96-well plate.

In the lung tissue analysis, the lung from each mouse was retrieved and weighed, followed by the separation of the left lung lobe, which was then stored at –80 °C until used for fungal load measurements. The right lung lobe was fixed in 10% buffered formalin and then subjected to a histopathological analysis. The remaining lung lobes were mechanically dissociated (BD Medimachine, Bruino, TO, Italy), resulting in cell suspensions that were filtered through a 70 µm cell strainer and then treated with the hemolytic buffer, when necessary. The cell-containing sediments obtained in the above steps were also distributed in the wells of a V-bottom 96-well plate and subjected to phenotyping. The liver and spleen were also retrieved and fixed in 10% buffered formalin, followed by routine processing for histopathology.

### Histopathological analysis and morphometry

Paraffin-embedded tissue samples were prepared routinely for histopathology. Tissue cuts were stained with hematoxylin–eosin (H&E), analyzed, and imaged under the DM LS (Leica, German) light microscope using the DFC420 image pick-up (Leica) and Leica Application Suite version 3.1.0 image program. Histological sections of lung fragments from each animal in both *Infected* and *Cy-Infected* groups were stained with H&E and then subjected to morphometric analysis. The lung perialveolar tissue area was determined from the digitized images in the morphometric analysis using the MetaMorph Microscopy Automation & Image Analysis software (Molecular Devices, California, USA). The hotspots in the pixels of the lesions of four animals from each experimental group were measured, followed by calculating the mean of each experimental group. The mean area of lesions was then determined and utilized to compare the microsporidian infection in animals between the two groups.

### Phenotypic characterization of immune cells

In order to block the Fc receptors of cells from Lung and BAL, these cells were incubated with the anti-CD16/CD32 antibody (diluted in PBS supplemented with 1% albumin-BSA) for 20 min. Afterward, the cells were washed and incubated with the following monoclonal antibodies: peridinin chlorophyll (PerCP)-conjugated or allophycocyanin (APC)-conjugated rat anti-mouse CD19, Phycoerithrin (PE)-conjugated rat anti-mouse CD23, Peridinin Chlorophyll Protein Complex (PerCP)-conjugated rat anti-mouse CD4, FITC-conjugated rat anti-mouse CD8, PE Cy7-conjugated rat anti-mouse F4/80, and pacific blue*-*conjugated rat anti-mouse CD11b (BD-Pharmingen, San Diego, CA). Next, the following cell phenotypes were identified in the evaluated cells: CD4^+^T cells (CD45^+^/CD4^+^), CD8^+^T cells (CD45^+^/CD8^+^), total macrophages (CD11b^+^F4/80^+^), interstitial macrophages (CD11b^+^F4/80^+^SIGLEC-F^-^), and alveolar macrophages (CD11b^-^F4/80^+^SIGLEC-F^+^). The cells were incubated at 4 °C, and after 30 min, the cells were washed, re-suspended in 200 µL of PBS, and evaluated using flow cytometry. The relevant data were acquired using BD Accuri^TM^ C6 (BD Biosciences, Mountain View, CA).

### Cytokine quantification

The plasma samples were thawed and evaluated for the levels of IL-2, IL-4, IL-6, IFN-γ, TNF-α, IL-17, and IL-10 using the BD CBA Mouse Th1/Th2/Th17 Cytokine Kit (BD Biosciences, CA, USA) according to the manufactures’ instructions. Briefly, 25 µL of each sample was added and allowed to capture the beads specific to the cytokines and PE (phycoerythrin)-labeled secondary antibodies. The sample was then incubated for 2 h at room temperature in the dark. Two-color flow cytometric analysis was performed using BD Accuri^TM^ C6 (BD Biosciences). The data were analyzed using FCAP Array analysis software version 3.0 (BD Biosciences, San Diego, CA, USA).

### Fungal load

The fungal load was determined by quantifying the fungal genomic DNA present in the lung tissue of each mouse using the real-time PCR (qPCR) technique. After euthanizing the mice, the lung samples were retrieved and weighed, followed by separating the left pulmonary lobe and freezing it in the RNA*later* ^®^ solution (Invitrogen™, USA) for a maximum of 30 days. Total DNA extraction was performed using the commercial DNeasy Blood & Tissue extraction kit (Qiagen^®^, Germany) according to the manufacturer’s instructions [12]. The extracted DNA was eluted using 100 µL of AE buffer and then stored at –20 °C until used for the qPCR analysis. The spores present in the lung tissue samples were quantified spectrometrically (BioDrop µLite, Biochrom) based on a standard curve generated based on the DNA extracted from the samples inoculated with 1 × 10^7^ purified spores/mL. Ten-fold serial dilutions of the standard, from 1 × 10^6^ to 1 × 10^1^ spores/µL, were performed in each amplification cycle (Cq) for the 18S rRNA gene of *E. cuniculi*. The best fit between the points was identified, and the coefficient of linear regression (R2) and the slope of the curve were calculated using the software. The quantitative polymerase chain reaction (qPCR) analysis was performed using the StepOne Plus^®^ Real-Time PCR (Applied Biosystems, Foster City, CA, USA). Amplification was performed in a total volume of 25 µL/reaction, which contained 12.5 µL of SsoAdvanced Universal SYBR^®^ Green Supermix 2X kit (Bio-Rad Laboratories, Inc., USA) along with 5 µL of the template DNA and the respective primers (20 pmol). Quality controls included positive DNA samples obtained from purified spores and non-template controls (NTC). The detection limit (LD) used in the present study was 10 spores/µL, the linear correlation coefficient (R2) was 0.998, and the amplification efficiency was 93.5%. The entire procedure was conducted in accordance with the Minimum Information Guide for Quantitative PCR Publication (MIQE) [13].

### Statistical analysis

T-test or variance analysis (ANOVA) was conducted followed by Tukey’s and Bonferroni’s multiple comparison post-tests for statistical analysis. All data were presented as means ±standard errors of the means (SEMs), using the significance threshold of α = 0.05 (p < 0.05). Graphs were plotted using “GraphPad Prism” version 5.0 for Windows ® (GraphPad Software Inc., La Jolla, CA, EUA).

## Results

### *E. cuniculi* oropharyngeal infection led to interstitial pneumonia associated with a mixed pattern of Th1, Th2, and Th17 cytokines

In the Sham group, the absence of lesions was predominant (Fig. 2A), although eventually, the desquamation of pneumocytes was observed, which was suggestive of broncho-aspiration pneumonia. This was attributed to the administration of sterile sodium chloride solution. The animals infected with *E. cuniculi* exhibited the presence of lymphoplasmacytic inflammatory infiltrates in the alveolar parenchyma (Fig. 2B and 2C). However, at times, a dense nodular mononuclear infiltrate was observed in the infected animals (Fig. 2D). In the microscopic examination, the presence of *E. cuniculi* spores inside cells was observed rarely. Leukocyte infiltrates were observed in the liver (Supplementary Fig. 1A and 1B), which was consistent with the presence of the pathogen, suggesting the spread of the infection acquired via the oropharyngeal route.

**Figure 2.**
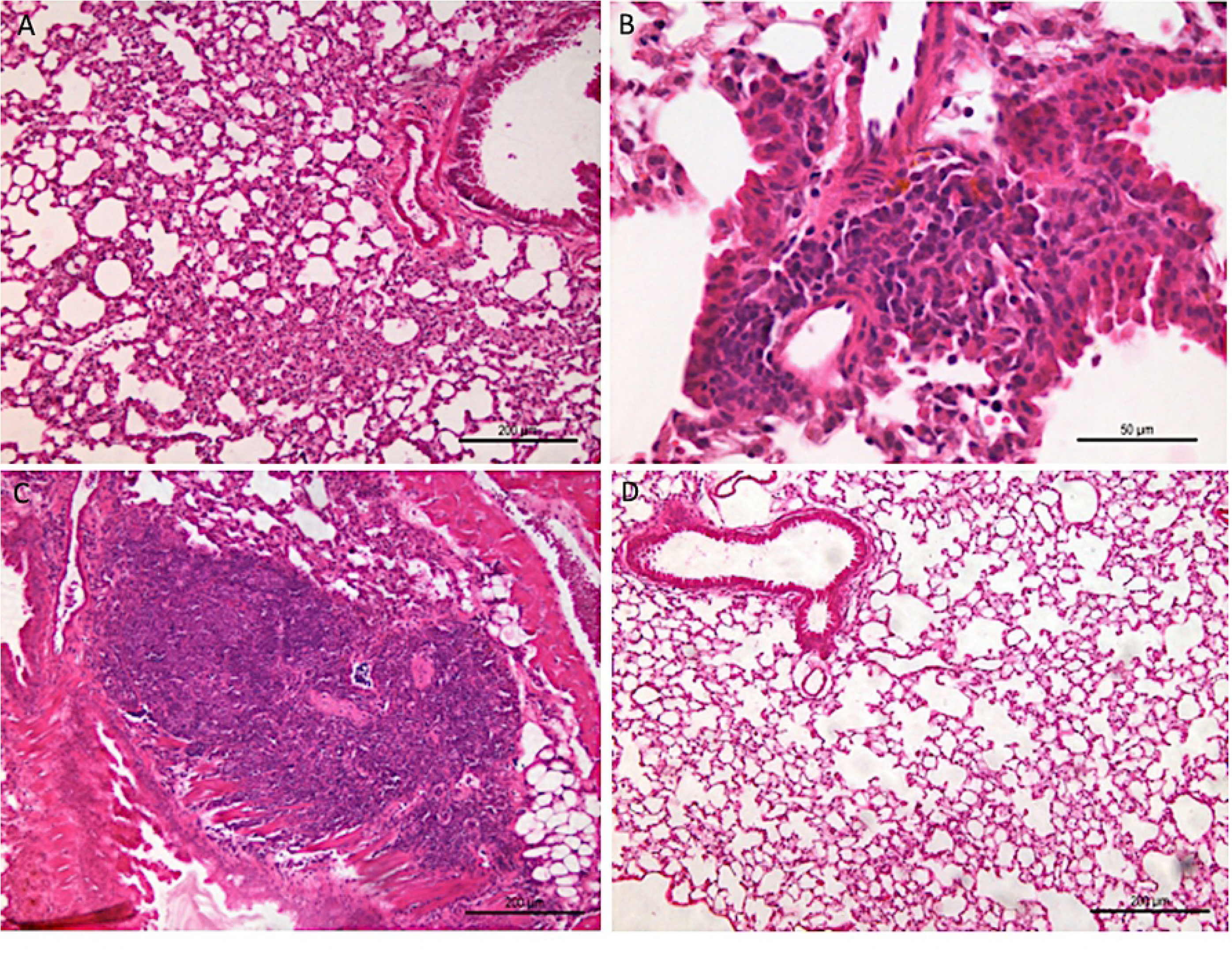
Photomicrographs of mice lungs infected with *E. cuniculi* spores via the oropharyngeal route or those administered with NaCl solution via the same route (Sham group). A) Sham group mouse without any signs of lesions in the lung parenchyma. B) Infected mice lung with diffuse thickening of alveolar walls. C) Dense mixed inflammatory infiltrate in the lung of the Infected group mice. D) Lung of mice infected with *E. cuniculi* exhibiting a predominantly mononuclear infiltrate with a dense nodular appearance (Hematoxylin and eosin [H&E] staining).

No difference in the percentages of CD8^+^ T lymphocytes was noted in the lung parenchyma and BAL between the Infected group mice and control mice. However, the percentage of CD4^+^ T cells in the lungs and that of B cells in the lungs and BAL were lower in the Infected group animals compared to controls (Fig. 3B and 3C). No differences in the levels of T and B lymphocytes in the blood were noted in all groups (Supplementary Fig. 2A).

**Figure 3.**
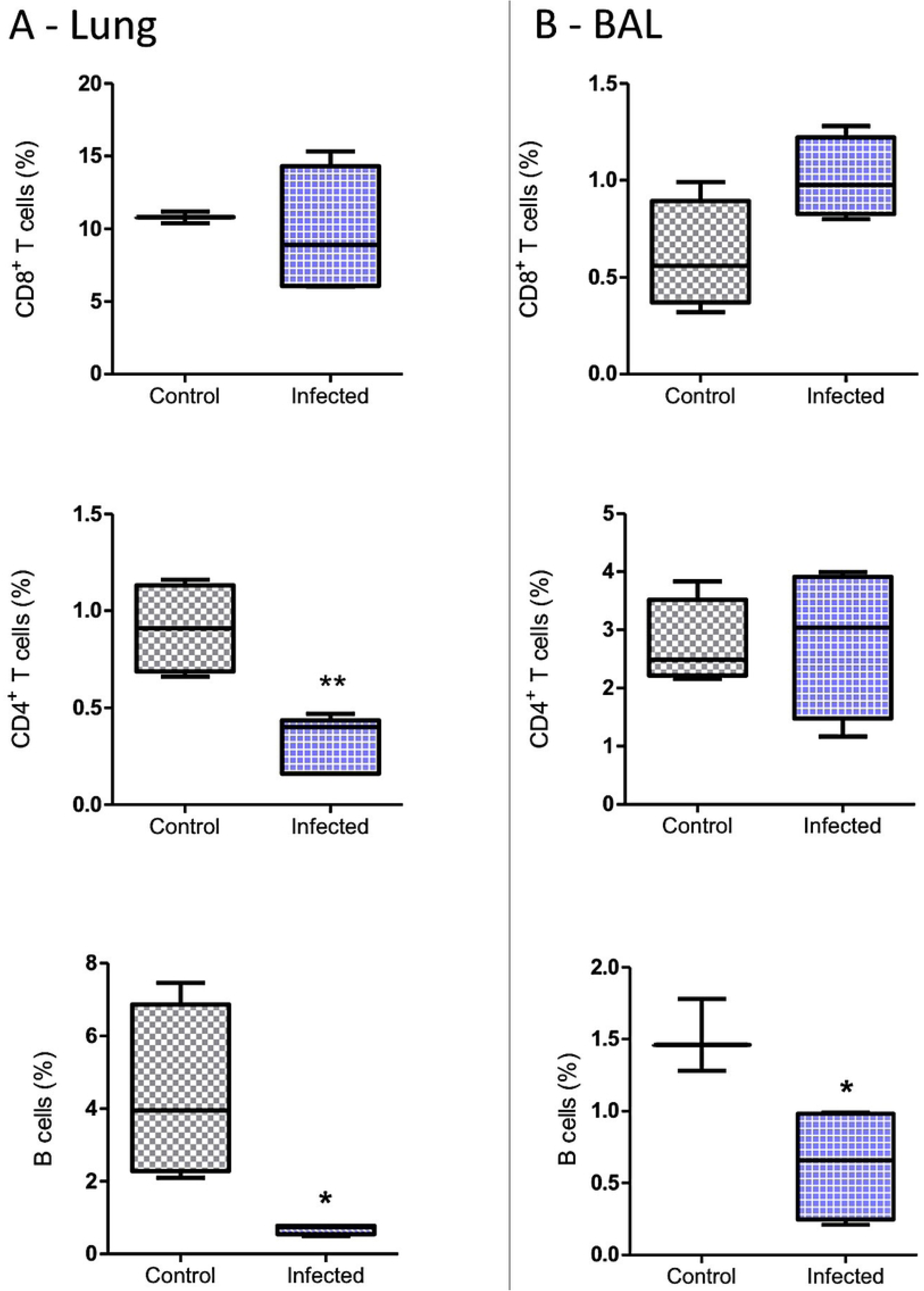
The total percentages (%) of CD8^+^, CD4^+^, T cell, and B cell populations in the lung and bronchoalveolar lavage (BAL) of the mice infected (Infected group) or not infected (Control) with *E. cuniculi* spores. (A) Lung. (B) BAL. The data presented are means ±standard errors of the means (SEMs) (*p < 0.05, **p < 0.01, T-test).

Further, the percentage of total macrophages was higher in the lungs of the Infected group of mice. This was attributed to the relative increase in interstitial macrophages (F4/80^+^CD11b^+^SiglecF^-^), while the percentage of alveolar macrophages (F4/80^+^CD11b^-^SiglecF^+^) decreased in this group (Fig. 4A). In BAL, the macrophage population was increased, which was attributed particularly to the higher percentage of alveolar macrophages (F4/80^+^CD11b^-^SiglecF^+^) compared to controls (Fig. 4B).

**Figure 4.**
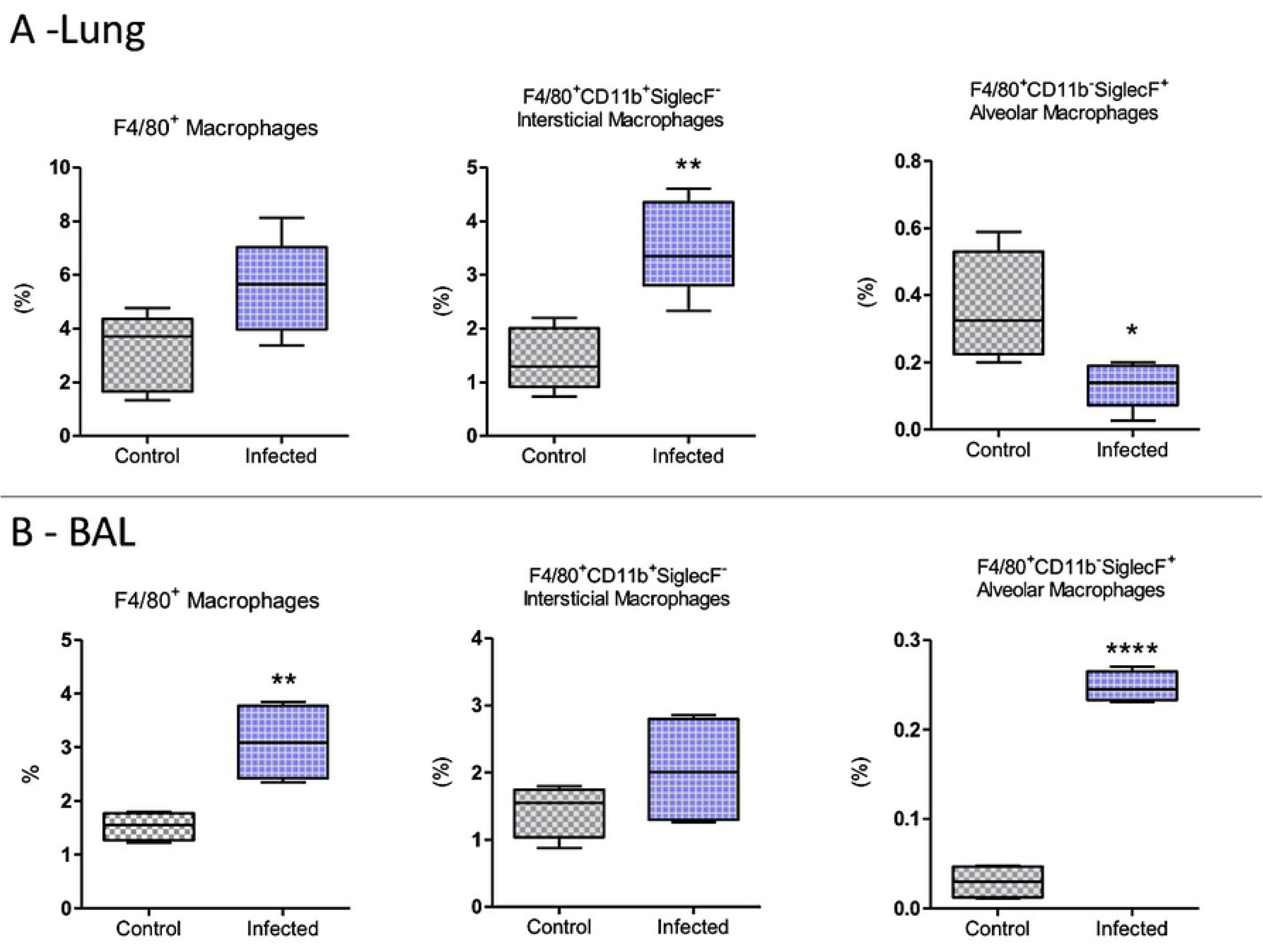
Macrophage populations in the lungs or bronchoalveolar lavage (BAL) of the mice infected (Infected group) or not infected (Control group) with *E. cuniculi* spores. A) Percentage of total macrophages (F4/80^+^), interstitial macrophages (F4/80^+^, CD11b^+^, and SiglecF^-^), and alveolar macrophages (F4/80^+^, CD11b^-^, and SiglecF^+^) in the lungs of mice. B) Percentage of total macrophages (F4/80^+^), interstitial macrophages (F4/80^+^, CD11b^+^, and SiglecF^-^), and alveolar macrophages (F4/80^+^, CD11b^-^, and SiglecF^+^) in BAL. The data presented are means ±standard errors of the means (SEMs) (*p < 0.05, **p < 0.01, T-test).

*E. cuniculi* infection led to increased plasma levels of pro-inflammatory and anti-inflammatory cytokines in the Th1, Th2, and Th17 profiles compared to controls (Fig. 5). In particular, levels of TNF-α, IL-2, and IL-6 increased 2 and 4 times relative to controls. Infection with *E. cuniculi* also resulted in increased IFN-γ levels. Among the anti-inflammatory cytokines, IL-4 exhibited increased levels compared to those in the control mice, while IL-10 was not detected. Interestingly, an increase was noted in the plasma IL-17 levels in the infected group compared to the Sham group, although no difference was noted compared to the control group (Fig. 5).

**Figure 5.**
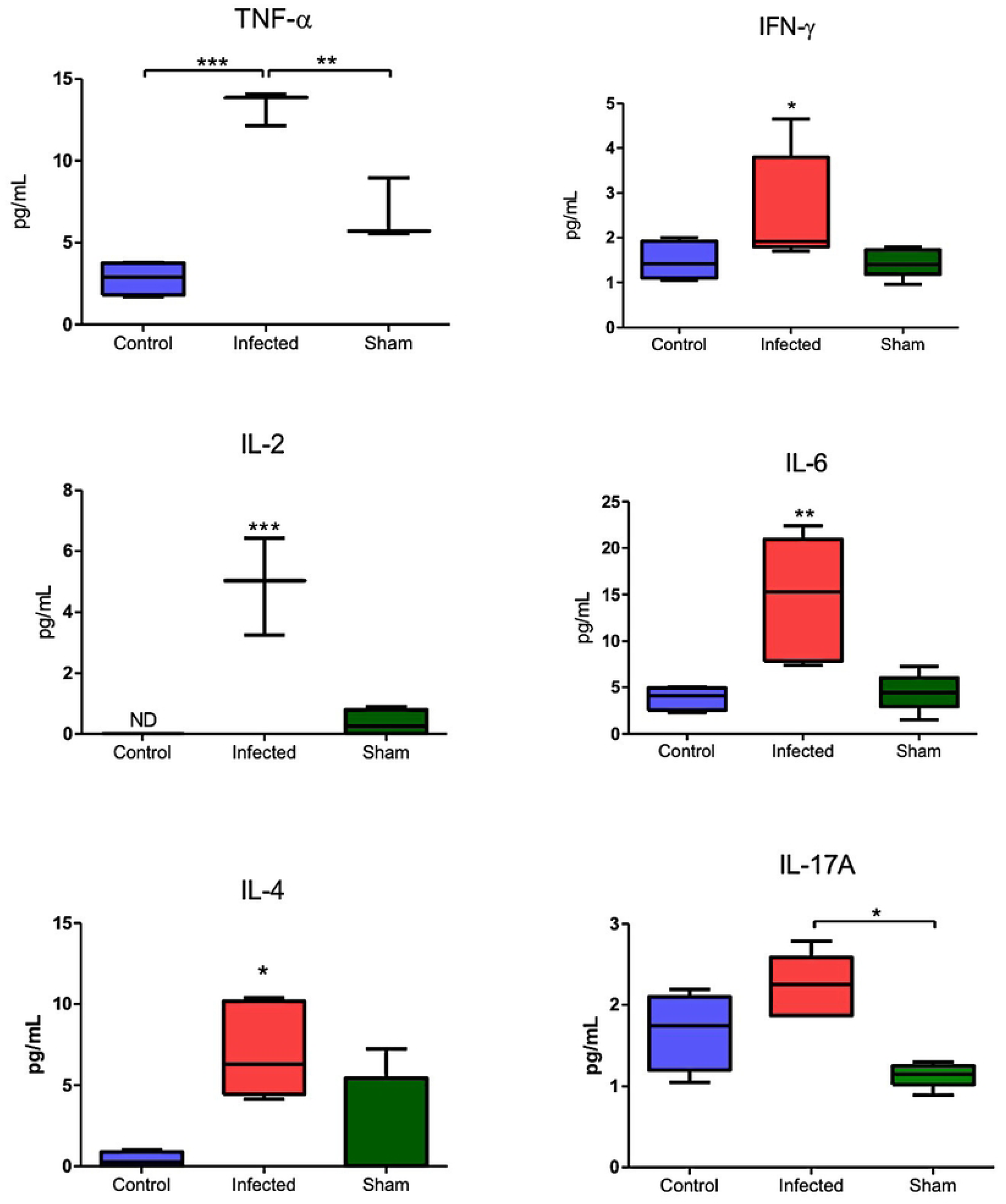
Plasma levels of TNF-α, IFN-γ, IL-6, IL-2, IL-17A, and IL-4 cytokines in the mice infected (Infected group) or not infected (Control group) with *E. cuniculi* spores and the uninfected mice that received NaCl solution via the oropharyngeal route (Sham). The data presented are means ±standard errors of the means (SEMs) [*p < 0.05, **p < 0.01, ***p < 0.001, one-way analysis of variance (ANOVA) along with multiple comparisons and Tukey’s post-tests].

### Reduction in fungal load was associated with increased CD8^+^ T lymphocyte levels in the lungs of animals immunosuppressed with Cy and infected with *E. cuniculi*

As an opportunistic pathogen, *E. cuniculi* leads to disseminated and severe infection in immunosuppressed individuals. Therefore, in the present study, mice that were previously treated with immunosuppressive doses of Cy were also infected with the pathogen via the oropharyngeal route for a comparative analysis of the infection model and the immune response. In the Cy-Infected group, lymphoplasmacytic inflammatory lesions were detected in the alveolar parenchyma, similar to the Infected group (Fig. 6A and 6B). However, when comparing the intensity of the lymphoplasmacytic inflammatory infiltrate by measuring the area of hotspots, the Infected group exhibited a significantly larger area compared to the Cy-Infected group (Fig. 6C). This finding is generally associated with the immunosuppressive and anti-inflammatory effects of Cy.

**Figure 6.**
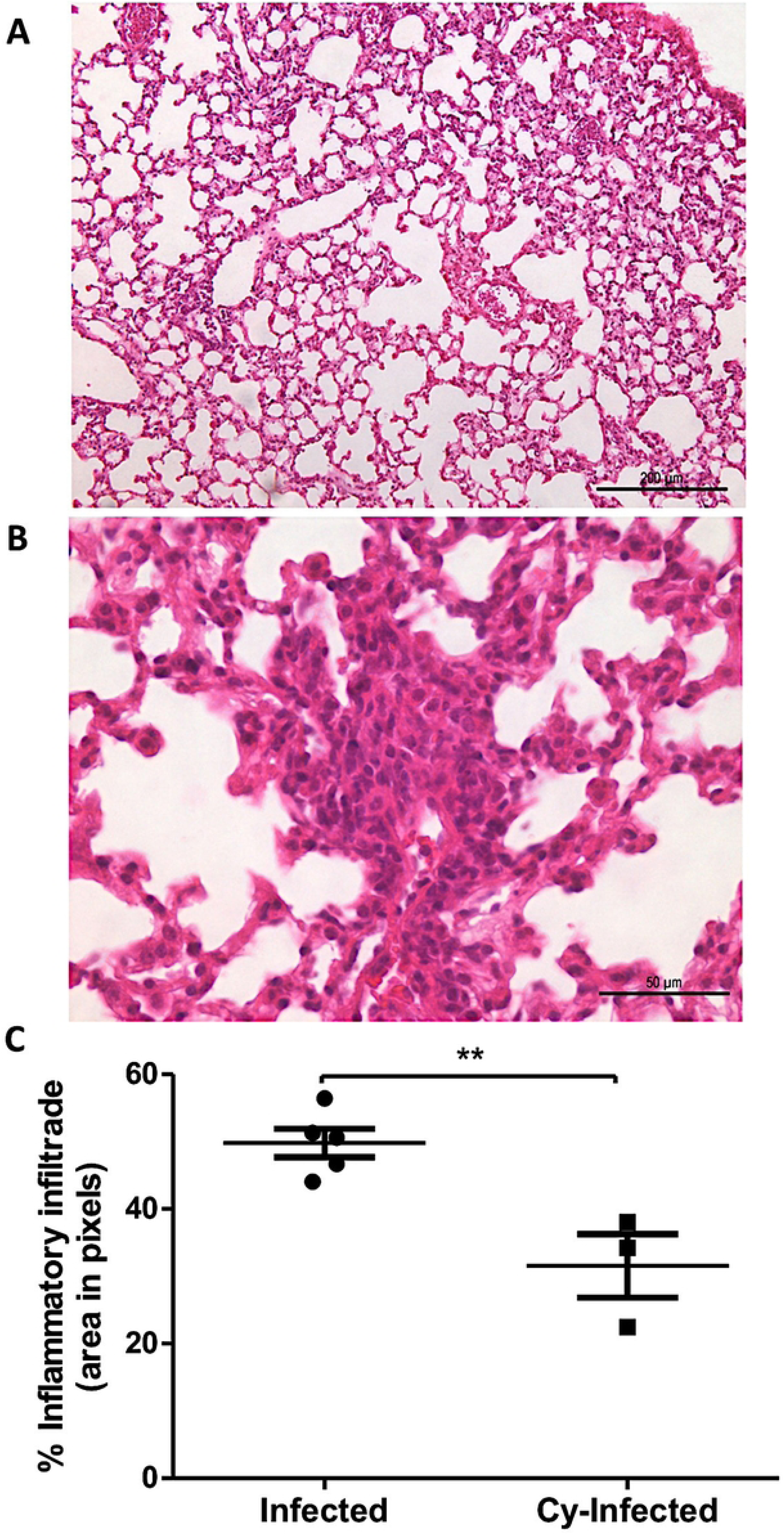
Histopathological lesions in the lungs of the mice treated with cyclophosphamide and then infected via the oropharyngeal route, and the comparative morphometric analysis of the inflammatory infiltrate in the mice infected with *E. cuniculi* and then treated [or not treated] with cyclophosphamide. A) Photomicrographs of the histological sections revealing discrete thickening of alveolar walls with diffuse distribution in the lungs of the mice treated with cyclophosphamide and then infected with *E. cuniculi* (Cy-Infected) [H&E]. B) Photomicrographs of the histological sections revealing a dense mononuclear infiltrate with a nodular shape in the lung of a Cy-Infected mouse [H&E]. C) Comparison of the percentage area of inflammatory infiltrate in the hotspots calculated using the metamorph software between the Infected and Cy-Infected groups. T-test revealed **p < 0.01.

The Infected and Cy-Infected groups were compared based on the quantification of fungal load in the mouse lung using qPCR. Surprisingly, the lung fungal load was higher in the Infected animals compared to the Cy-Infected animals (Fig. 7). However, despite the immunosuppression induced using cyclophosphamide, the percentage of CD8^+^ T lymphocytes in the lungs of the Cy-Infected group was higher compared to the Cy-Uninfected group (Fig. 8A). Meanwhile, the CD4^+^ T lymphocytes were increased in the BAL of Cy-Infected group relative to the Cy-Uninfected group, while the percentage of CD8^+^ T lymphocytes decreased in the Cy-Infected group (Fig. 8B). No differences were noted in the levels of blood T and B lymphocytes in all groups (Supplementary Fig. 2B). Microsporidiosis plays a critical cytotoxic role, due to which this marked difference in the CD8 T lymphocyte population could explain the lower fungal load noted in the Cy-Infected group despite the immunosuppressive effect of Cy.

**Figure 7.**
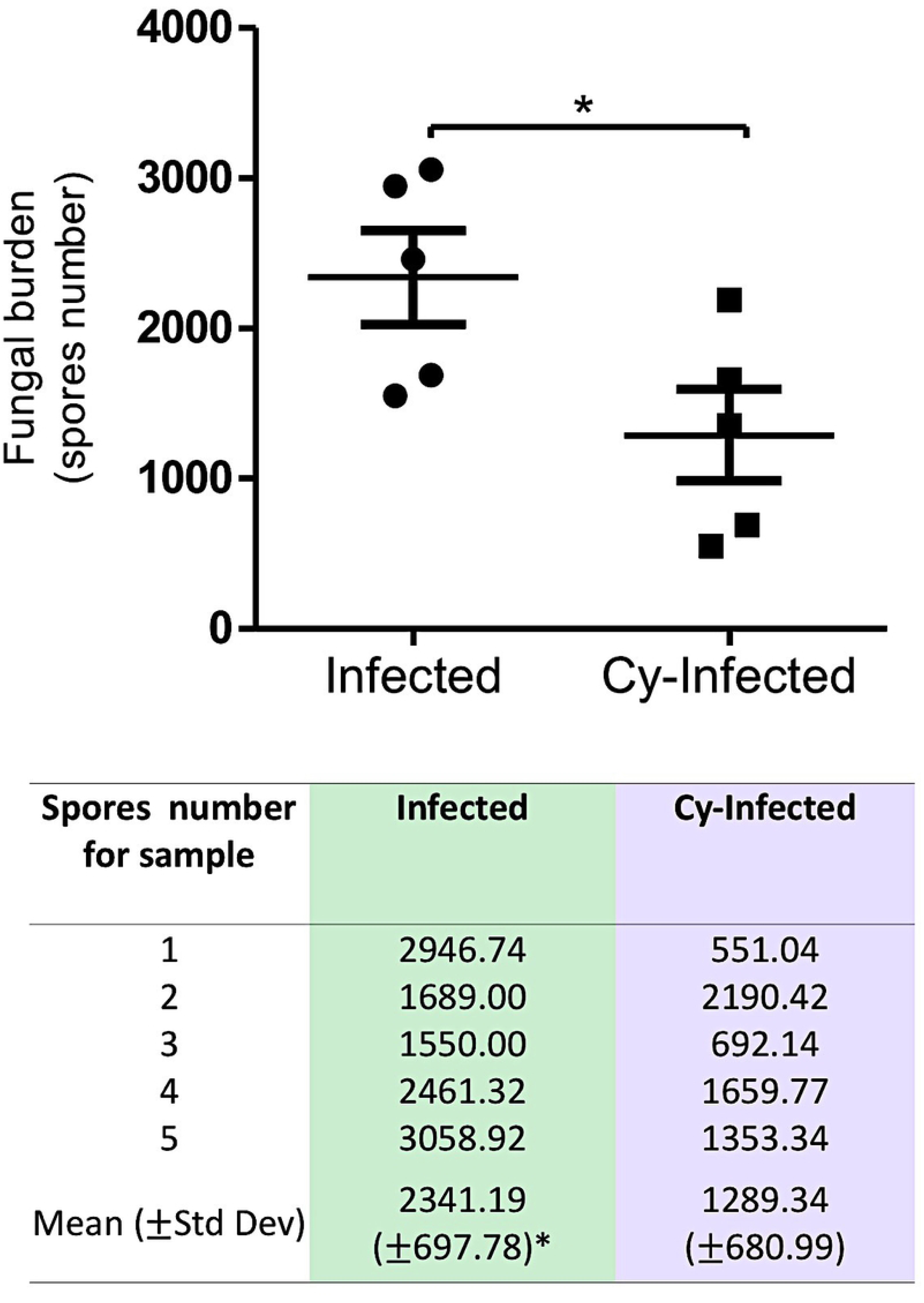
Quantification of fungal load in the lung parenchyma of the mice infected with *E. cuniculi* spores (Infected group) or immunosuppressed with cyclophosphamide (Cy) and then infected (Cy-Infected group) using the real-time PCR (qPCR) technique (Student’s *t*-test, *p* < 0.05*).

**Figure 8.**
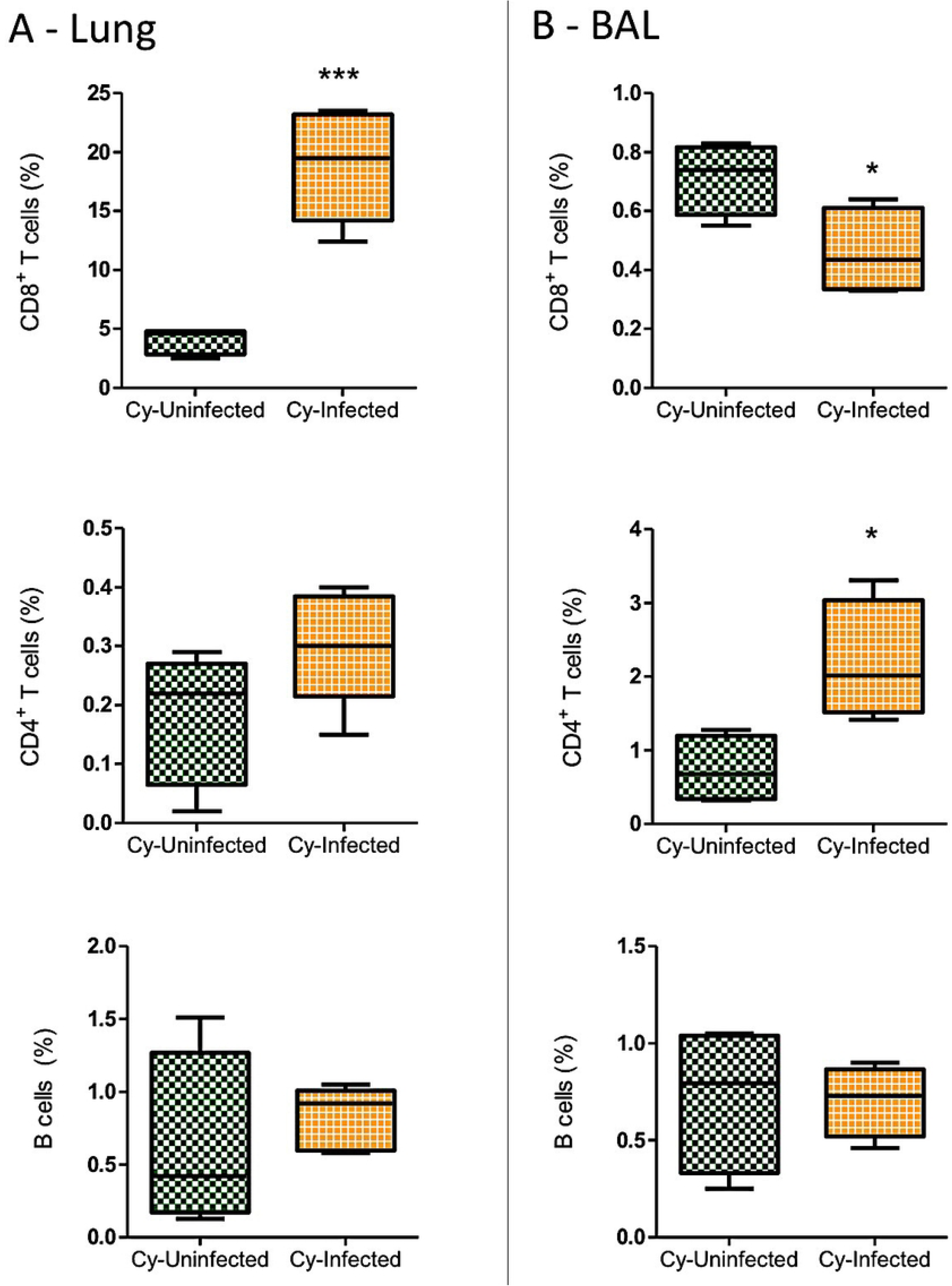
The total percentage (%) of CD8^+^ and CD4^+^ T cell and B cell populations in the lung and bronchoalveolar lavage (BAL) of the mice immunosuppressed with cyclophosphamide (Cy) and then infected (Cy-Infected group) or not infected (Cy-Uninfected group) with *E. cuniculi* spores. (A) Lung. (B) BAL. The data presented are means ±standard errors of the mean (SEMs) (*p < 0.05, **p < 0.01, T-test).

The percentage of macrophages in the lung did not differ between the Cy-Infected and Cy-Uninfected groups, despite a trend toward reduced levels of alveolar macrophages (Fig. 9A). However, in BAL, the percentage increase in the macrophages in the Cy-Infected group was evident, particularly due to increased levels of interstitial macrophages (F4/80^+^CD11b^+^SiglecF^-^) (Fig. 9B). In the blood, increased levels of macrophages were detected in Infected, Cy-Infected, and Cy-Uninfected groups (Supplementary Fig. 3).

**Figure 9.**
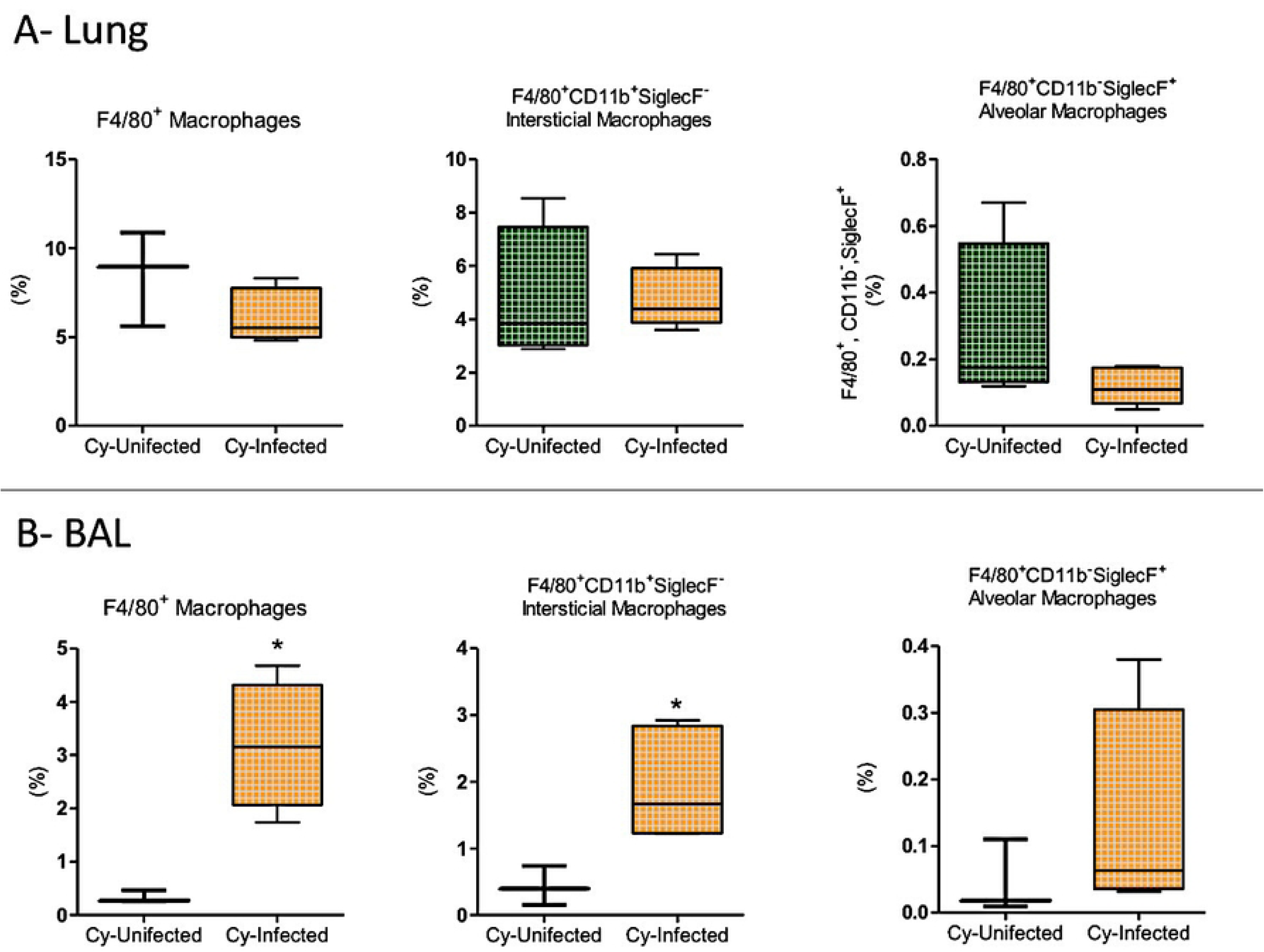
Macrophage populations in the lungs or bronchoalveolar lavage (BAL) of the mice immunosuppressed with cyclophosphamide (Cy) and then infected (Cy-Infected group) or not infected (Cy-Uninfected group) with *E. cuniculi* spores. A) Percentage of total macrophages (F4/80^+^), interstitial macrophages (F4/80^+^, CD11b^+^, and SiglecF^-^), and alveolar macrophages (F4/80^+^, CD11b^-^, and SiglecF^+^) in the lungs of mice. B) Percentage of total macrophages (F4/80^+^), interstitial macrophages (F4/80^+^, CD11b^+^, and SiglecF^-^), and alveolar macrophages (F4/80^+^, CD11b^-^, and SiglecF^+^) in BAL. The data presented are means ±standard errors of the mean (SEMs) (*p < 0.05, **p < 0.01, T-test).

The cytokine profiles determined for the blood samples from the animals treated with Cy were quite different, with decreased levels of inflammatory cytokines TNF-α, IFN-γ, and IL-2 noted in the Cy-Infected group compared to the other groups, except for just IL-6, which was increased in this group (Fig. 10). In addition, the anti-inflammatory cytokines IL-4 and IL-17A were increased in the Cy-Infected group (Fig. 10). According to these results, the animals in this group presented a less inflamed profile.

**Figure 10.**
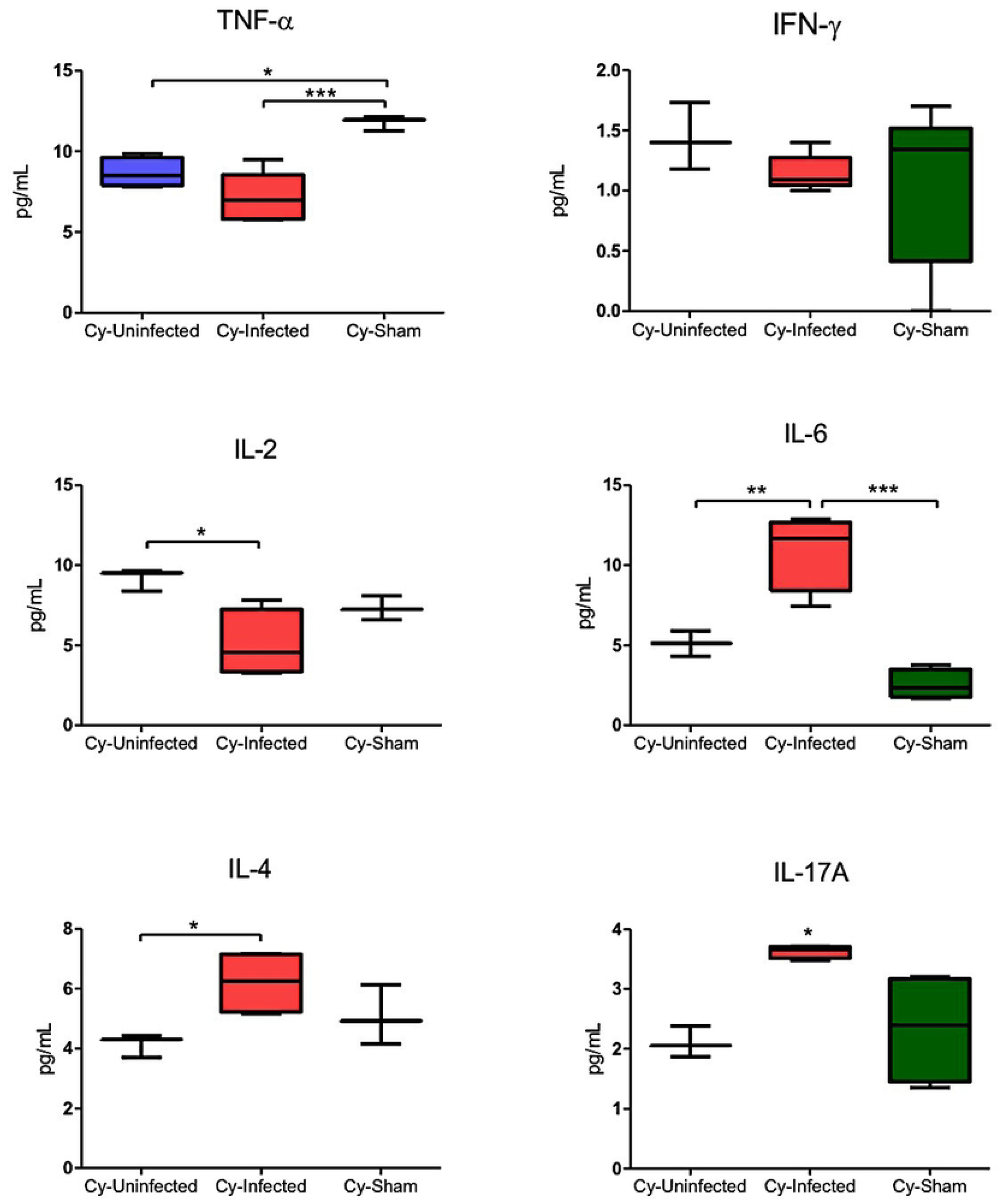
Plasma levels of TNF-α, IFN-γ, IL-6, IL-2, IL-17A, and IL-4 cytokines in the mice immunosuppressed with cyclophosphamide (Cy) and then infected (Cy-Infected group) or not infected (Cy-Uninfected group) with *E. cuniculi* spores and the uninfected and Cy-immunosuppressed mice that received the NaCl solution via the oropharyngeal route (Cy-Sham group). The data presented are means ±standard errors of the mean (SEMs) [*p < 0.05, **p < 0.01, ***p < 0.001; one-way analysis of variance (ANOVA) along with multiple comparisons and Tukey’s post-tests].

The greater resistance to the pathogen observed in the Cy-Infected group was attributed to the higher percentage of CD8^+^ T lymphocytes detected in the lungs of the animals in this group. In addition, the Infected group mice exhibited a further inflamed cytokine profile, with increased levels of TNF, IFN, IL-2, and IL-6. These results demonstrated that infection via the oropharyngeal route facilitated the establishment of *E. cuniculi* infection in the lung tissues of mice with a consequent inflammatory response, resulting in predominant interstitial pneumonia, with the non-immunosuppressed animals being less resistant to the infection compared to Cy-Infected immunosuppressed animals.

## Discussion

The present study demonstrated the occurrence of interstitial pneumonia in all mice infected with *E. cuniculi* via the oropharyngeal route, thereby filling a gap in the epidemiological knowledge of this zoonosis, as this route of infection had not been demonstrated experimentally. The occurrence of pneumonia characterized by lymphoplasmacytic and occasionally mononuclear inflammatory infiltrate observed in the present study corroborated the results obtained in other previous studies on the intraperitoneal route of infection. In a previous study, for instance, *E. cuniculi* led to variable degrees of degeneration, necrosis, and desquamation of the bronchiolar epithelium and inflammatory infiltrates comprising mononuclear cells in the lung parenchyma of animals [14]. However, the pneumonia observed in the present study was more severe and the infection was further intense in non-immunosuppressed animals, which is different from the findings of the aforementioned authors who reported a greater number of lesions caused by *E. cuniculi* in immunosuppressed mice [14]. Cyclophosphamide is a non-specific cell cycle drug based on an alkylating agent derived from nitrogen mustard. It works by incorporating an alkyl group into the DNA, thereby preventing DNA synthesis and division. The immunosuppressive effects of cyclophosphamide (Cy) are attributed to its action on the DNA within the inflammatory cells, in addition to its suppressive effect on the bone marrow. The cytotoxic effect of this drug is due to the interaction of its alkylating metabolites with DNA. Treatment with Cy reportedly aggravated *E. cuniculi* infection, rendering it more severe, widespread, and with significantly increased lethality [10, 15, 16, 17]. Contrary to the observations related to the intravenous infection of mice with *E. cuniculi* [18], in the present study, the fungal load was lower in the immunosuppressed group compared to the mice not treated with cyclophosphamide and then infected with *E. cuniculi*, which is quite intriguing. This difference may be attributed to the high population of CD8^+^ T lymphocytes detected in the lungs of infected groups of mice. The Cy-Infected group exhibited a higher percentage of CD8^+^ T lymphocytes compared to the other groups. Previous studies have demonstrated that CD8^+^ T lymphocytes are the effector cells in encephalitozoonosis [19], and this could explain the lower fungal load noted in this group in the present study.

The lungs have two distinct populations of macrophages: alveolar macrophages (AM), which are in close contact with the type I and II epithelial cells of the alveoli, and interstitial macrophages (IM), which are located in the parenchyma between the microvascular endothelium and the alveolar epithelium. The functional phenotype of AM is modulated by the unique microenvironment of the lung, which comprises intimate contact with the epithelial cells, high oxygen tension, and exposure to surfactant-rich fluid, all of which are critical for tissue homeostasis, host defense, elimination of surfactant and cellular debris, pathogen recognition, initiation and resolution of lung inflammation, and repair of damaged tissue [20]. In the present study, an increase in MA populations (F4/80^+^CD11b^-^ SiglecF^+^) was noted in the lungs and BAL of non-immunosuppressed infected animals, along with a greater fungal load and inflammatory infiltrate, indicating a further pronounced *Encephalitozoon* pneumonia. A milder pneumonia was observed in the mice deficient in MAs and infected with human metapneumovirus, with lower body weight loss, lung inflammation, and airway obstruction noted compared to the mice carrying MAs. On the other hand, pneumonia caused by syncytial virus was reported to be more severe in MA-deficient mice, suggesting that MA activity might vary depending on the causal agent [21]. The presence of AMs, reported in the present study, could be associated with further severe pneumonia in the encephalitozoonosis established via the oropharyngeal route.

MAs, as the first line of defense, initiate the innate immune response in the lung. However, 2 MA phenotypes may exist: the classically activated macrophages (MA–M1) and the alternatively activated macrophages (MA–M2). M1 macrophages respond to microbial factors and Th1 pro-inflammatory cytokines, thereby leading to glycolytic metabolism, which is associated with the release of inflammatory cytokines, increased bacterial killing, and recruitment of immune cells into the lung parenchyma and alveolus. M2 macrophages, on the other hand, are induced, by exposure to Th2 cytokines, leading to oxidative metabolism, which is associated with the release of anti-inflammatory cytokines, phagocytosis of apoptotic cells (efferocytosis), and collagen deposition, which contribute to the resolution of inflammation and repair of damaged tissues [20].

In the present study, it was hypothesized that the increased levels of MAs associated with the mixed cytokine profile of Th1 (IFN-γ, TNF-α, and IL-6) and Th2 (IL-4) observed in the serum samples from animals could be indicative of the concomitant presence of MAs from the M1 and M2 profiles in non-immunosuppressed animals. In a previous study, efferocytosis was observed to be favorable to *E. cuniculi* and induced the profile of M2 macrophages, thereby favoring the multiplication and dissemination of the pathogen [22], similar to the change in the profile of macrophages noted in the animals infected with *E. cuniculi*. In this context, AMs could likely be contributing to the greater susceptibility of animals to the pathogen, acting as a “Trojan Horse”.

Interstitial macrophages (IMs) constitute the second line of defense in the lungs against invading pathogens. However, it is important to highlight that a dichotomy exists between IMs. Schyns et al. [23] demonstrated that the bronchial population of CD206^+^ IMs has a superior capacity to secrete immunoregulatory cytokines, including IL-10, consistent with the reported homeostatic functions of regulating inflammation and fibrosis. On the other hand, CD206^−^ MIs exhibited a typical antigen-presenting cell profile and were demonstrated to regulate the T cell-related processes. The increase in the levels of MIs in the lungs of infected mice observed in the present study and its association with a further severe infection might be suggestive of the adoption of an immunoregulatory profile, although further investigation is necessary for clarification. The levels of MA and MI macrophages were, on the other hand, not increased in the lungs of Cy-treated and infected mice, which demonstrated the resolution of infection during the period evaluated and also an increased population of CD8^+^ T lymphocytes. These inferences, however, have to be understood better.

Previous studies have demonstrated that increased levels of Th1 cytokines in the serum of animals are associated with an increase in reactive oxygen and nitrogen products, which are important factors regarding resistance to *Encephalitozoon* infections in mice [19, 24, 25, 26]. In the present study as well, a predominance of Th1 cytokines (TNF-α, IFN-γ, IL-2, and IL-6) was noted in the serum of infected mice.

IL-6 is a pleiotropic cytokine with several biological functions, including the regulation of the immune system through the production of other anti-inflammatory and pro-inflammatory cytokines [27]. Mice with diabetes mellitus (DM) exhibit greater susceptibility to encephalitozoonosis, which manifests as higher levels of IL-6 [19]. This was also demonstrated in the findings of the present study for animals not treated with Cy. In contrast, Cy-treated animals exhibited elevated IL-6 levels associated with the resolution of infection, suggesting an anti-inflammatory role of IL-6 and reiterating the ambiguity of this cytokine.

Th17 cytokines are categorized into six IL-17 subtypes, from A to F, which are produced by a broad spectrum of cell populations, including γδT, NKT, CD8^+^ T cells, neutrophils, microglia, and mast cells [4, 28, 29]. Other important sources of IL-17 include myeloid cells (e.g., in the kidneys and lungs) and Paneth cells in intestinal crypts [28]. IL-17A responses are required to control the infections caused by fungal pathogens, including *Aspergillus fumigatus, Pneumocystis carinii, Cryptococcus neoformans*, and *Candida albicans* [30, 31, 32, 33, 34, 35]. IL-17C, on the other hand, induces lethal inflammation by exacerbating the secretion of pro-inflammatory cytokines, thereby contributing to the development of systemic fungal infection [33]. These paradoxical effects have to be explored comprehensively.

Under the conditions described in the present study, the increase in the serum IL-17 levels noted in the Cy-Infected group, which also had the lowest inflammation and fungal load, suggests that the role of this cytokine is consistent with the results obtained in previous studies on other fungal infections [30, 31, 32, 33, 34, 35].

## Conclusion

The oropharyngeal route of infection promoted *E. cuniculi* pneumonia in the mice treated or not treated with cyclophosphamide, with a greater severity noted in non-immunosuppressed mice. Therefore, this model of infection could be established as a suitable model of interstitial pneumonia via aspiration, with the potential for the identification of the role of lung immune cells in infections caused by intracellular pathogens such as *E. cuniculi*.

---

**Supplementary Figure 1.** Hepatic granuloma (A and B) in the mice infected with *E. cuniculi* spores and then treated with cyclophosphamide (H&E).

**Supplementary Figure 2.** The total percentage (%) of CD8^+^ and CD4^+^ T cell and B cell populations in the blood. A) The percentage of CD8^+^ and CD4^+^ T cell and B cell populations in the mice infected (Infected group) or not infected (Control group) with *E. cuniculi* spores. B) The percentage of CD8^+^ and CD4^+^ T cell and B cell populations in the mice immunosuppressed with cyclophosphamide (Cy) and then infected (Cy-Infected group) or not infected (Cy-Uninfected group) with *E. cuniculi* spores. The data presented are means ±standard errors of the mean (SEMs) (T-test).

**Supplementary Figure 3.** The absolute number or percentage (%) of macrophage populations in the blood. A) Mice infected (Infected group) or not infected (Control group) with *E. cuniculi* spores. B) Mice immunosuppressed with cyclophosphamide (Cy) and then infected (Cy-Infected group) or not infected (Cy-Uninfected group) with *E. cuniculi* spores. The data presented are means ±standard errors of the mean (SEMs) (*p < 0.05, ****p < 0.0001, T-test).

